# Transcriptional coregulation *in cis* around a contact insulation site revealed by single-molecule microscopy

**DOI:** 10.1101/2025.03.04.641491

**Authors:** Maciej A. Kerlin, Ilham Aboulfath-Ladid, Julia Roensch, Chloé Jaubert, Aude Battistella, Kyra J. E. Borgman, Antoine Coulon

## Abstract

Chromosome conformation in mammals is closely related to gene regulation. Within topologically associating domains, where genomic contacts are enriched, genes tend to show correlated expression across tissues and conditions, suggesting domain-wide mechanisms coregulating multiple genes, such as enhancer sharing or local histone mark spreading. At the single-cell level, where transcription occurs in sporadic bursts, transcriptional coordination has been observed between proximal genes, but how the local folding of mammalian chromosomes influences gene coregulation *in cis* at individual alleles remains unclear. Using single-molecule microscopy, we imaged nascent transcription from three adjacent genes located around a strong contact insulation site at the *FOS* locus, during the estrogen response in human breast cancer cells. To interpret this data, we developed two new analysis approaches to dissect the sources of (co)variation in gene activities: one to separate allele-extrinsic, allele-intrinsic, and gene-autonomous components; and another one to quantify the contributions of burst co-occurrence and burst size correlations. We find that transcriptional variability is largely gene-autonomous, yet correlations between genes display distinct patterns and occur almost exclusively *in cis*. Correlations are stronger between proximal and less insulated genes. However, unexpectedly, substantial correlations also occur across the strong insulation site and, under certain conditions, two proximal genes on the same side can exhibit uncorrelated burst occurrences. By disentangling burst co-occurrence from burst size correlations, we reveal transcriptional patterns suggesting two distinct coregulatory mechanisms influenced by local chromosome folding.

## Introduction

The mammalian genome is highly organized in both space and sequence. At the sub-megabase scale, chromosomes fold into domains of enriched contacts, called topologically associating domains^1,2^. These domains have been shown to constrain gene regulation by enhancers^3,4^ and have been proposed to function as insulated regulatory units^5,6^, though some evidence nuances this view^7,8^. Still, the tendency of genes within these domains to show correlated expression across tissues and conditions^1,9–11^–especially when functionally related^12^– suggests the existence of domain-wide coregulatory mechanisms, such as enhancer sharing or histone mark spreading.

Transcriptional correlations provide a powerful tool to study gene coregulation at different scales, from population-level analyses to single-cell studies^13,14^. At the single-cell level, where transcription occurs in sporadic bursts, proximity between genes has long been linked to transcriptional correlations^15^. More recently, correlation patterns between genes have been used to investigate local regulatory mechanisms in mammals^16,17^, flies^18,19^, and yeast^20,21^. However, how the local folding of mammalian chromosomes shapes gene coregulation *in cis* at individual alleles remains unclear.

Here, we combine single-molecule imaging with new data analysis approaches to investigate transcriptional correlation patterns among three adjacent human genes flanking a strong contact insulation site. We show that transcriptional coordination within a contact domain occurs at the level of individual alleles. However, we also uncover unexpected features that challenge a strict role for genome folding in transcriptional coordination and reveal correlation patterns suggesting two distinct coregulatory mechanisms, whose origins we speculate on.

## Results and Discussion

### Population-level data identify local, contact-dependent correlations in the transcriptional response of genes to estrogen

We first used public GRO-seq data^22^ to quantify, genome-wide, the local correlations between the (population-averaged) transcriptional response of genes to estradiol (E2) in MCF7 cells. We found that genes within a few hundred kilobases tend to have similar temporal profiles of transcriptional response (Figure 1A), unless strongly insulated from one another as measured by Hi-C^23^ (Figure 1B). This suggests local coregulatory mechanisms linked to 3D organization (itself shaped by genomic distance and chromosome folding mechanisms), consistent with our current understanding of enhancer-gene control^3^. Yet, it remains unclear whether such coregulation between estrogen-responsive genes occurs locally *in cis* and how it manifests through transcriptional bursting.

**Figure 1.**
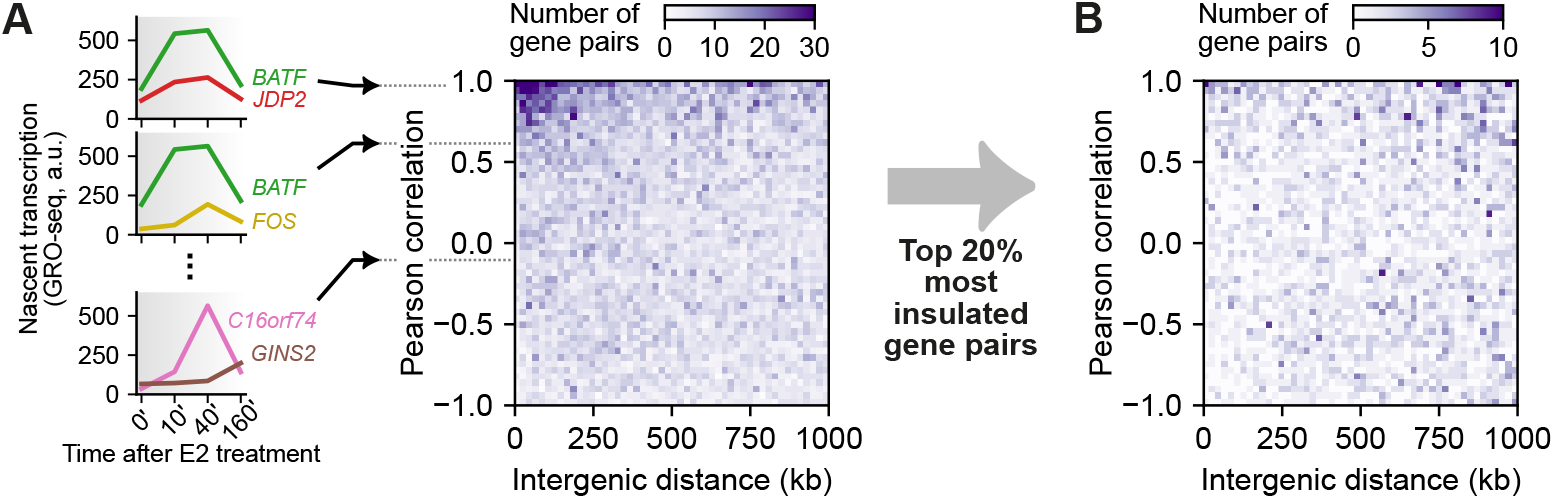
Population-level data suggest local transcriptional coordination in response to estrogen. (**A**) From GRO-seq measurements^22^ of nascent transcription over time in response to estradiol (E2) in MCF7 cells, the similarity of transcriptional time profiles between genes is assessed using the Pearson correlation coefficient (i.e. insensitive to differences in offset or scale of the time profiles) and plotted against their genomic distance as a two-dimensional histogram. (**B**) Same analysis using only gene pairs separated by a strong contact insulation site as measured by Hi-C^23^.

### Single-molecule imaging of the joint activities of three genes at individual alleles with combined DNA-RNA FISH

To address this, we chose three adjacent genes: the proto-oncogene *FOS*, separated from *JDP2* and *BATF* by a strong contact insulation site (Figures 2A, S1A). All three genes are upregulated following E2 exposure, while showing basal transcriptional activity in the absence of E2 (Figures 1A, S1B). The genomic proximity of these genes (< 250 kb), their strong functional relationships (AP-1 family transcription factors^24^), and the abundance of estrogen receptor α (ERα) binding sites in the vicinity (Figure S1B) hints towards the existence of local mechanisms that coregulate them.

**Figure 2.**
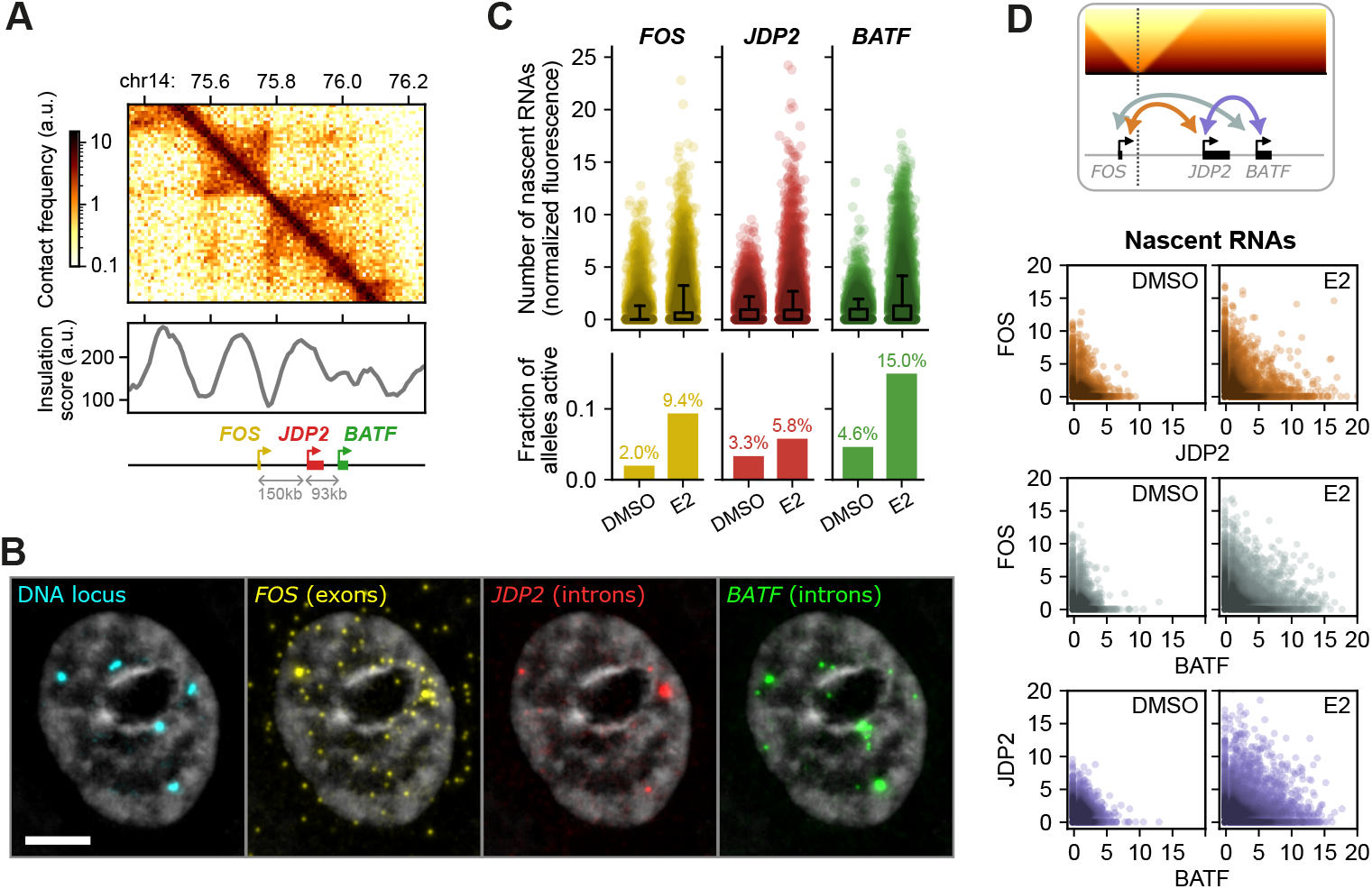
Measuring nascent transcription of three adjacent genes at individual alleles. (**A**) Hi-C contact map and insulation score (low values indicate strong contact insulation) at our genomic locus of interest. (**B**) MCF7 cell imaged in five colors after combined oligo-based DNA FISH and single-molecule RNA FISH (gray: DAPI). Five allele loci are visible in the DNA channel. In the RNA channels, bright spots colocalize with allele DNA loci, revealing ongoing bursts of nascent RNAs, while dim spots correspond to single RNAs (see Methods and Figure S1D-I). Shown are maximum-intensity projections of band-pass filtered images. Scale bar: 5 μm. **(C)** Quantification of nascent transcription of *FOS, JDP2*, and *BATF*, measured by FISH at individual alleles, following a 40-min treatment with 100 nM estradiol (E2) or vehicle control (DMSO), represented as number of nascent RNAs (top) and fraction of active alleles (bottom). (**D**) Same data, represented as joint distributions of number of nascent RNAs at individual alleles, for all three gene pairs, in uninduced (DMSO) and induced (E2) conditions.

We combined single-molecule RNA FISH with oligonucleotide-based DNA FISH to co-image the transcriptional activities of *FOS, JDP2*, and *BATF*, at each allele of the locus in MCF7 cells (Figure 2B). With an oligonucleotide library^25^ spanning the whole locus, we used the DNA FISH channel to identify individual alleles (which copy number substantially varied between nuclei; Figure S1D-E). We observed bright RNA FISH spots at the identified DNA loci, corresponding to transcription sites, from which we estimated the number of nascent RNAs using the brightness of single RNAs as a reference (Figures S1F-G). With this approach, we calculated the distributions of nascent RNA counts and the fractions of active alleles for each gene following treatment with E2 or vehicle control (DMSO) for 40 min (Figure 2C), while maintaining cells in G1 using palbociclib (Figure S1C). We confirmed that all three genes go from basal to upregulated transcription upon E2 exposure (Figure 2C) and undergo transcriptional bursting (Figure S1H). The resulting joint distributions of nascent RNAs at individual alleles for all three gene pairs (Figure 2D) are the basis for analyzing how transcription is locally coordinated across the locus.

### Two mathematical approaches to separate sources of (co)variations in gene activity

Transcriptional correlations between genes were broken down into multiple additive components using two strategies (Figures 3A-B and S2).

**Figure 3.**
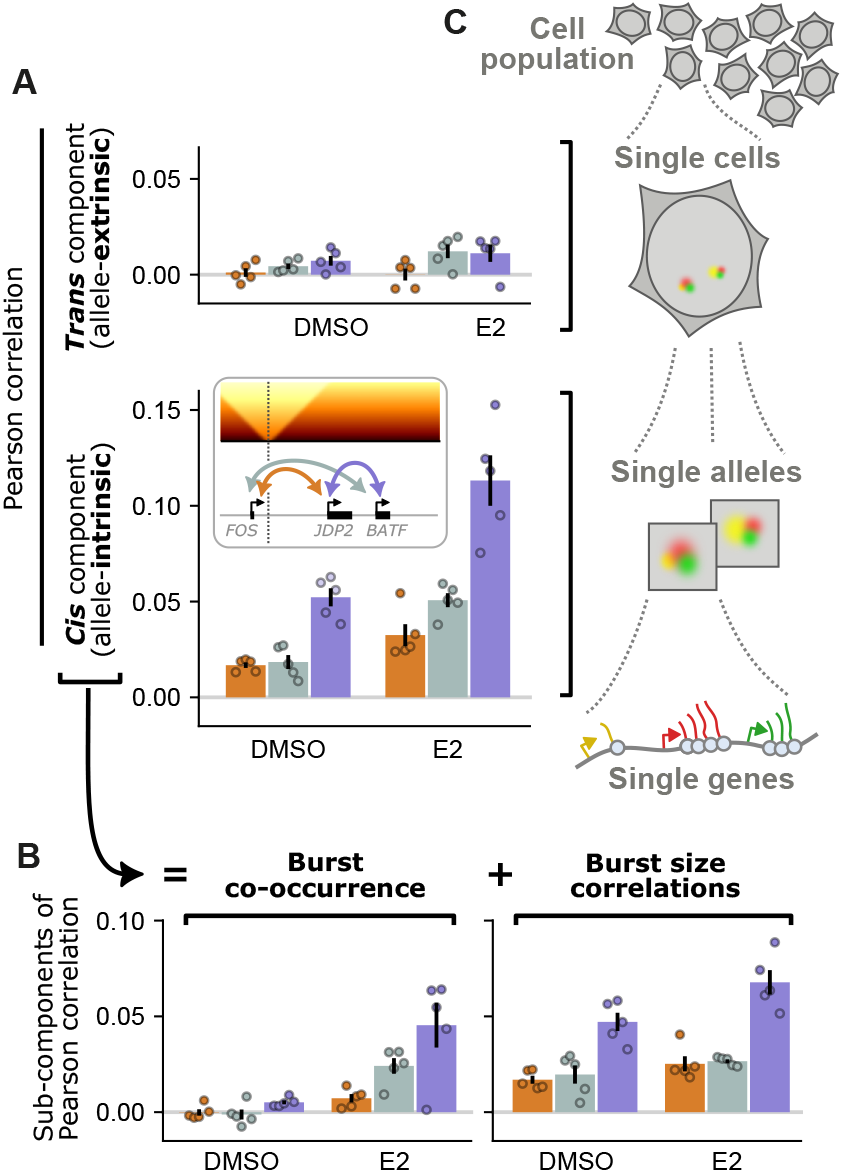
Multilevel decomposition of transcriptional correlations. (**A**) For each gene pairs, the Pearson correlation coefficient between nascent RNAs at individual alleles is separated into two additive components, isolating the contribution of *trans* and *cis* effects. (**B**) The latter is further separated into two additive components, quantifying correlations due to burst co-occurrence and burst size correlations, respectively. Averages of 5 replicates (dots) are shown as bars, with standard errors as error bars. See Methods for statistical tests and *p*-values. (**C**) Our data and computational approaches yield a multiscale understanding of transcriptional correlation patterns, from population-level heterogeneity to allele-specific coupling and gene bursting dynamics.

First, we developed an approach to separate out global effects due to cell-to-cell variability (e.g. cell cycle, cell-wise differences in E2 responsiveness…). Our method extends the classical *intrinsic*/*extrinsic noise* framework^26^, generalizing it to variable allele copy numbers and multiple genes per allele, yielding *allele-intrinsic* and *allele-extrinsic* pairwise correlations between genes (Figure 3A). It relies on computing covariance matrices from both allele-level measurements and their cell-wise sums, then using arithmetic to extract covariances across all possible cell-wise *trans* permutations and isolate *cis*-level effects—i.e. what would be observed in the absence of cell-to-cell variability (see Methods; Figure S2A-B). While applied here between *single genes* at *individual alleles* within *single cells* of a *population* (Figure 3C), our generic framework can apply recursively —e.g. from single-cell and/or spatial-omics data (Figure S2C-D)— opening avenues for multiscale analyses of heterogeneities and correlation patterns^14^.

Second, to further dissect transcriptional correlations, we reasoned that, just as bursting is characterized by burst frequency (how often bursts occur) and burst size (number of RNAs per burst), correlations between genes may arise from burst co-occurrence (e.g. partially synchronized burst events, or simply covarying burst frequencies) and/or burst size correlations (e.g. correlated burst magnitudes, even if occurrence may be uncorrelated). To separate these effects, we combine the *law of total covariance* with a perturbation approach to break down the covariances into several components: one stemming from the contingency table of the burst/no-burst binarized data, while the rest characterizes correlations involving burst sizes (Figure S2E).

### Transcriptional fluctuations are predominantly gene-autonomous

Before interpreting the transcriptional correlation patterns between *FOS, JDP2*, and *BATF*, an important observation is that these correlations occur almost exclusively *in cis* and are rather moderate (Figure 2A). Indeed, the transcriptional variability of each gene considered individually is dominated by allele-intrinsic –or *cis*– effects (with only minor contributions from allele-extrinsic –or *trans*– effects; Figure S3) and even *in cis*, the covariances between genes remain moderate compared to the variances (Figure S3). Together, these findings indicate that, in our experiments (i.e. on populations of cells exposed to the same E2 treatment and arrested in G1), transcriptional fluctuations are largely gene-autonomous.

### The most proximal and least insulated gene pair exhibits the strongest *cis* correlations

Transcriptional correlations *in cis*, although moderate, offer a basis to study local mechanisms that coregulate transcription. Across all three gene pairs and in both uninduced (DMSO) and induced (E2) conditions, we find that the strongest *cis* correlation occurs in the most proximal and least insulated gene pair (*JDP2*-*BATF*; Figure 3A). This observation, made here for the first time at the single-allele level, aligns with our population-level analysis (Figure 1) and with previous reports at the population^1,9^ and single-cell^12^ levels, and support the possibility of domain-wide *cis-*coregulatory mechanisms relying on 3D chromosome organization (e.g. enhancer sharing, contact-dependent histone mark spreading).

In addition, while we cannot disentangle the contributions of 1D genomic distance and 3D contacts (since the least insulated genes are also the closest), correlation strength does not appear to be strictly determined by gene order nor by genomic distance, in line with earlier observations at larger scales^17^.

### Transcriptional correlation patterns nuance the role of local genomic contacts in gene coregulation

Although our results and many observations in the field^1,3,12,17^ suggest 3D chromosome folding as a dominant factor in gene (co)regulation, two unexpected observations nuance this view.

First, we find that substantial transcriptional correlations can occur across a strong contact insulation site. Specifically, correlations between genes on either side of the contact insulation site (*FOS*-*JDP2* and *FOS*-*BATF*) are only ∼3-fold weaker than for genes on the same side (*JDP2*-*BATF*) (Figure 3B), despite the ∼7:1 intra-to-inter-domain contact ratio between the two domains delimited by the insulation site (Figure SA1). This indicates that contact insulation does not make genes operate in isolation, but rather only moderately uncouple their activities, consistent with the idea that regulatory mechanisms may bypass contact insulation sites^7^.

Conversely, the two genes on the same side of the contact insulation site (*JDP2*-*BATF*) can, under certain conditions, show a surprising lack of transcriptional coordination. Indeed, separating the contributions of burst co-occurrence and burst size correlations reveals that the former is negligible in uninduced (DMSO) conditions for all three gene pairs (Figure 3B). Among other implications discussed below, this finding shows that *JDP2* and *BATF*, two adjacent and proximal genes (< 100 kb apart) without a strong contact insulation site between them, can exhibit mostly uncorrelated burst occurrences.

### Burst occurrence and size statistics outline two distinct mechanisms of local gene coregulation

Transcriptional correlations *in cis* are present in uninduced conditions and increase upon E2 induction (Figure 3A). Our analysis disentangling burst co-occurrence from burst size correlations reveals a remarkably clear explanation for this behavior: burst co-occurrence arises almost exclusively upon E2 induction, while burst size correlations account for all basal pre-induction correlations and remain largely unchanged by induction (Figure 3B). This suggests two distinct coregulatory mechanisms, both of which appear shaped by local chromosome folding (strongest correlations in the most proximal and least insulated gene pair; Figure 3B).

The molecular origins of these two modes of coregulation remain speculative. Enhancers having been shown to modulate burst frequency^3,14,16^ and to coordinate burst timing in flies^18,19^, we propose that the burst co-occurrence we observe, which is almost entirely induction-dependent, may stem from the activity of the many ERα-bound enhancers at the locus (Figure S1A). This contrasts with an earlier report of disfavored co-bursting between genes sharing a single enhancer^16^, possibly reflecting here a collective behavior of many enhancers. In turn, burst size correlations reveal preexisting allele-to-allele variations that remain mostly unchanged during the 40 min of E2 induction. Burst sizes having been associated to histone marks, we propose that the burst size correlation we observed may originate from contact-dependent local spreading of histone marks^27,28^, hence constrained by genome folding and contact insulation patterns.

Further work will be needed to elucidate the mechanisms underlying such local transcriptional coordination between genes.

**Figure S1.**
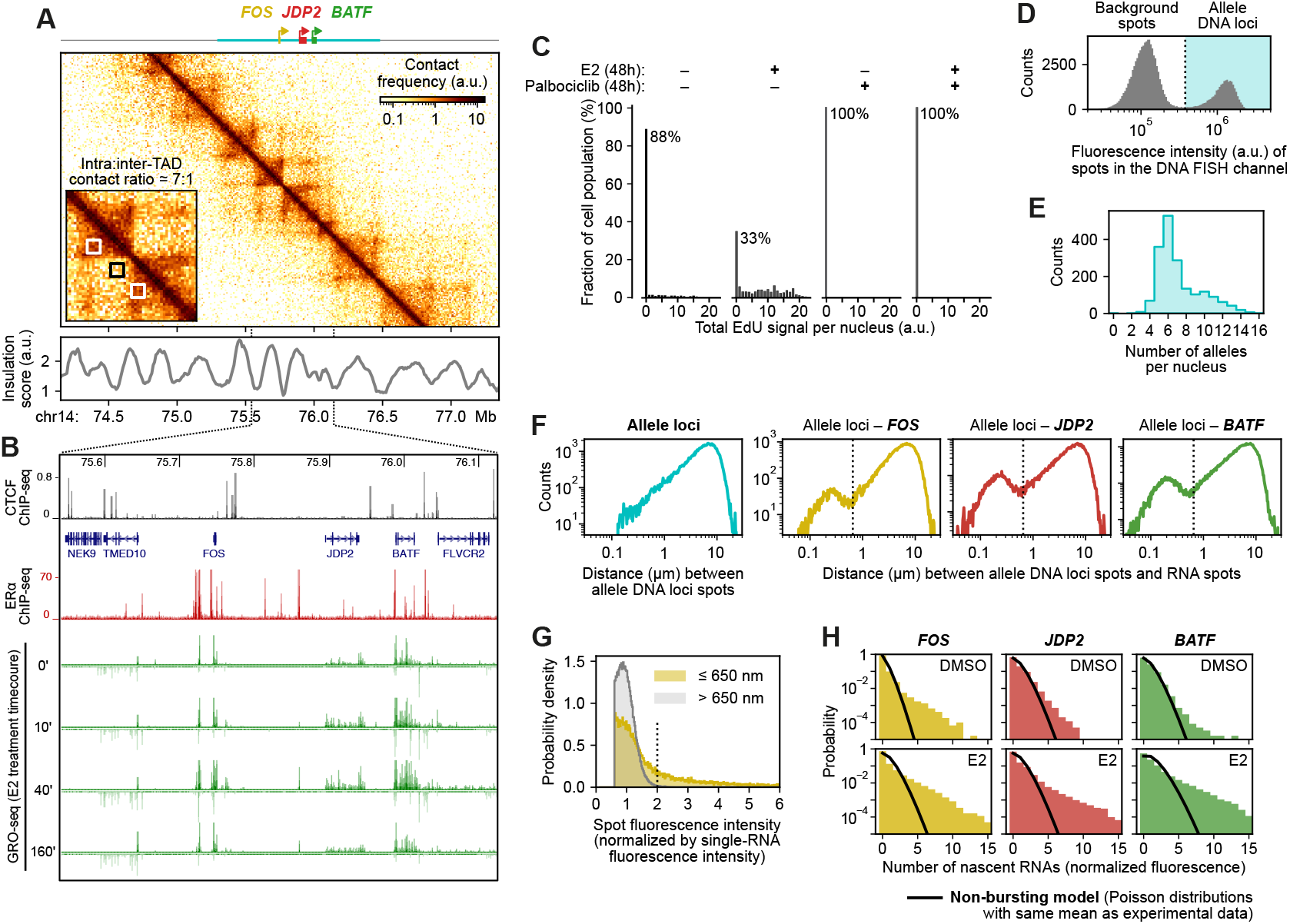
Local genomic context and transcription at the *FOS*-*JDP2*-*BATF* locus. Genomic context of our locus of interest described by publicly available data from MCF7 cells: **(A)** Hi-C contact map^23^ and an insulation score (low score indicates strong insulation), **(B)** ChIP-seq for CTCF and ERα on E2 treated cells^29,30^, and GRO-seq over time after E2 treatment^22^. **(C)** Histograms of EdU signal measured in MCF7 cells, after a 48 hrs treatment with different drug combinations, followed by 8 hrs of EdU incorporation. Percentages of arrested cells are indicated. **(D)** Distribution of fluorescence intensity of spots detected within nuclei of MCF7 cell, in the DNA FISH channel, using 12,005 fluorescent oligonucleotide probes labeling our locus of interest. Vertical line: intensity threshold above which a spot is considered to be an allele locus. **(E)** Histogram of the number of allele loci per nucleus. **(F)** Distributions of pairwise distances (left) between allele loci or (right) between allele loci and RNA FISH spots in each of the three RNA FISH channels (*FOS, JDP2*, and *BATF*). Vertical line: 650 nm. **(G)** Distributions of fluorescence intensity of *FOS* RNA spots, separating spots within 650 nm of an allele locus (yellow) from the rest (gray). The latter being considered single RNAs, the peak of their distribution is used, here and thereafter, to normalize fluorescence intensities. Vertical line: intensity threshold for an RNA spot to be unambiguously brighter than a single RNA, used to classify alleles a transcriptionally active. **(H)** Distributions of the number of nascent RNAs at individual alleles for all three genes, in uninduced and induced conditions. Black curve: expected distributions if the genes were not undergoing transcriptional bursting, i.e. producing individual RNAs as random and uncorrelated events.

**Figure S2.**
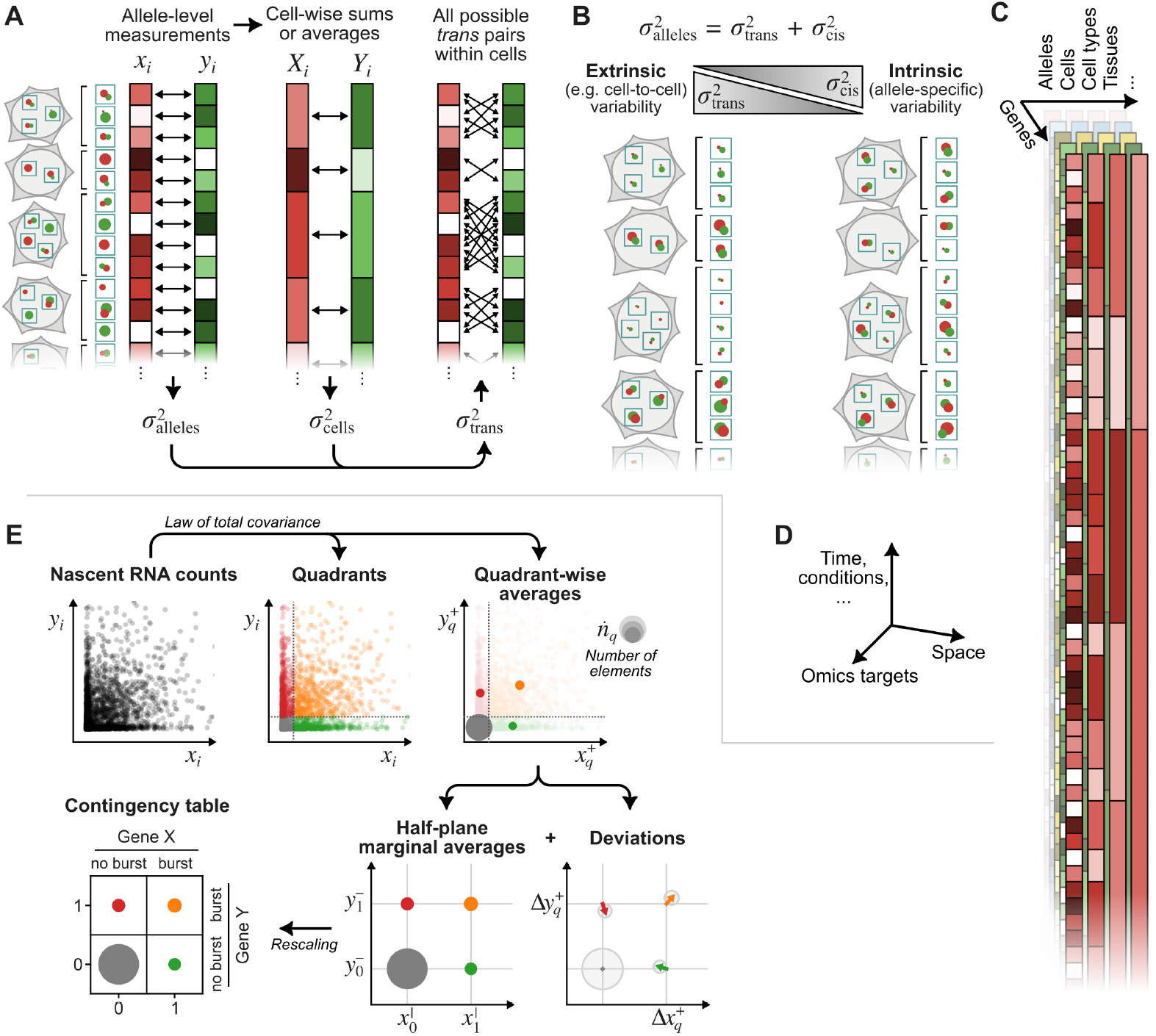
Mathematical approaches to decompose transcriptional correlations. (**A**) Considering measurements of nascent RNAs on two genes at individual alleles in single cells, the covariance 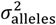 of such measurements and the covariance 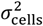 of their sums or averages within cells can be combined to deduce the covariance 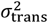 across all the possible *trans* permutations. Intuitively, 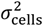 contains the products of the cell-wise sums (i.e. *X*_1_*Y*_1_ + *X*_2_*Y*_2_ +…) which, when expanded in terms of single-allele measurement (e.g. (*x*_1_ + *x*_2_ + *x*_3_)(*y*_1_ + *y*_2_ + *y*_3_) + (*x*_4_ + *x*_5_)(*y*_4_ + *y*_5_) +…), contains all intra-cellular pairs (i.e. ***x***_**1**_***y***_**1**_ + *x*_1_*y*_2_ + *x*_1_*y*_3_ + *x*_2_*y*_1_ + ***x***_**2**_***y***_**2**_ + *x*_2_*y*_3_ + *x*_3_*y*_1_ + *x*_3_*y*_2_ + ***x***_**3**_***y***_**3**_ + ***x***_**4**_***y***_**4**_ + *x*_4_*y*_5_ + *x*_5_*y*_5_ + ***x***_**5**_***y***_**5**_ +…), out of which the ***cis*** terms can be subtracted out using 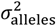 (i.e. containing only the ***cis*** terms, ***x***_**1**_***y***_**1**_ + ***x***_**2**_***y***_**2**_ + ***x***_**3**_***y***_**3**_ + ***x***_**4**_***y***_**4**_ + ***x***_**5**_***y***_**5**_ +…). See Methods. (**B**) The *trans* covariance 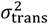 captures extrinsic factors (cell-to-cell differences in cell cycle, availability of machinery…) and equals the allele-level covariance 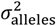 under the null hypothesis that genes X and Y at the same alleles do not correlate more or less than any other (non-*cis*) pair within cells. The residual 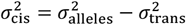 hence captures intrinsic variability, i.e. effects that are specifically occurring in *cis*. Illustrated are the two extreme scenarios where the variability is dominated by extrinsic factors (left, i.e. 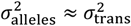) or by intrinsic factors (right, i.e. 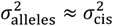). (**C**) Our general approach can be used recursively on data sets with several hierarchical levels: the cell-wise aggregated data (sums or averages of allele-level data) can itself be aggregate further at the cell-type level, then at the tissue levels… etc, each time decomposing further the origin of the variability. (**D**) Potential application in a broader context includes single-cell and/or spatial multi-omics data, which may include time (development, response to stimuli…) and experimental conditions, as well as other ‘omics’ measurements than gene transcription (e.g. scATAC, scCUT&Tag, …). (**E**) The joint distribution of nascent RNA counts from two genes can separated into four quadrants, indicating for each allele whether either gene is bursting or not. The total covariance of RNA counts can be written as the sum of the covariance within quadrants and the covariance of the average RNA counts within quadrants, which can themselves be expressed as a rescaled version of the contingency table and deviation terms. Separating out the contribution of the contingency table from the covariance of RNA counts characterizes burst co-occurrence, while all the other terms involve burst sizes.

**Figure S3.**
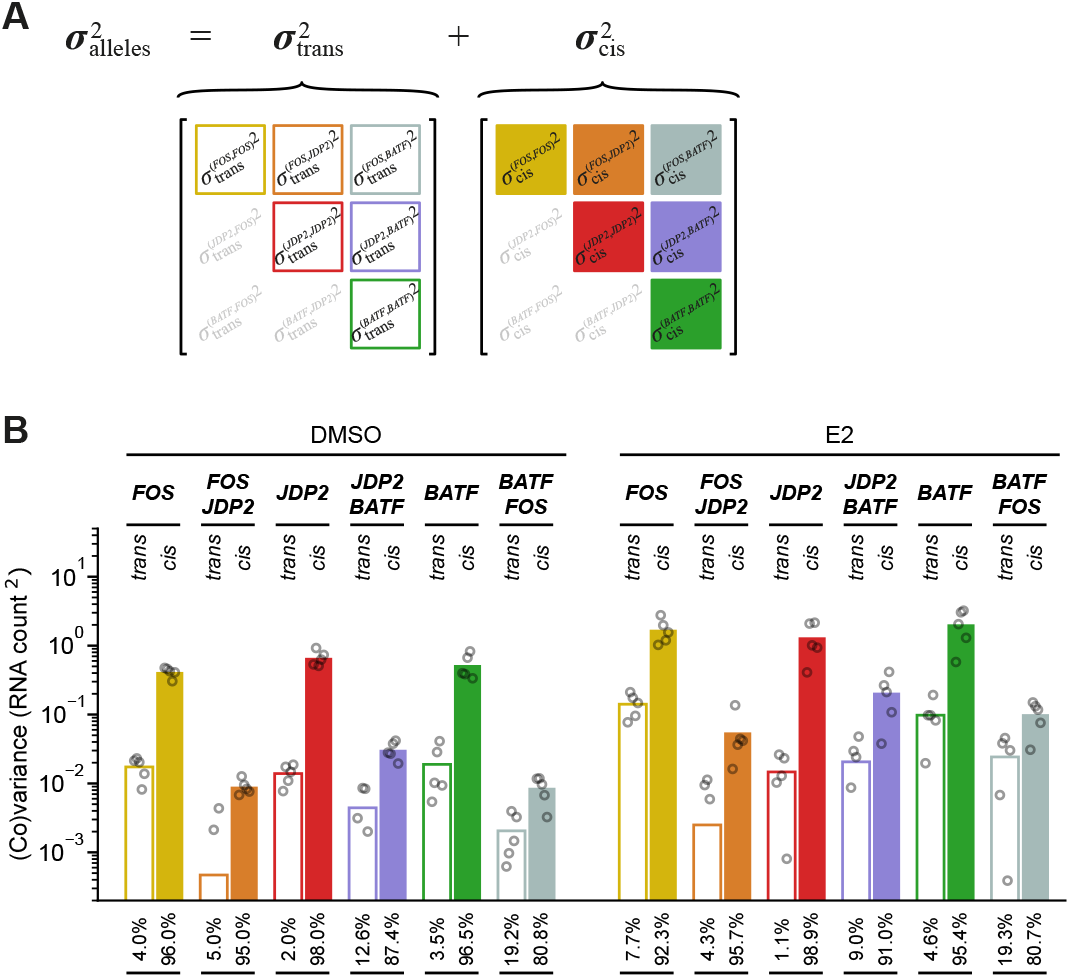
Covariance matrices. **(A)** The covariances calculated across all the alleles measured from a cell population can be separated into two components, respectively capturing *trans* effects (i.e. cell-to-cell heterogeneity) and *cis* effects (i.e. (co)variations that are not explained by cell-wise effects). See Methods and Figure S2A-B. Each component corresponds to a covariance matrix, the values of which are shown in panel (B) with the same color code. **(B)** *Trans* and *cis* components of the variances and covariances of nascent RNA counts, in uninduced (DMSO) and induced (E2) conditions. Each bar is the average of n = 5 replicates, shown as points. Numbers below the bars indicate the contribution of the *cis* and *trans* to the total (co)variance 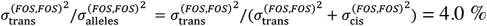 in DMSO conditions).

## Material and Methods

### Cell lines and cell culture

Human breast cancer cells MCF7 (ATCC) were grown at 37°C with 5% CO_2_ in DMEM medium (Gibco, ref. 41966052) supplemented with 10% FBS (Gibco, ref. 10500064) and 5% Penicillin/Streptomycin (thereafter referred to as *regular culture medium*) and split twice a week.

For FISH sample preparation, 200k cells were plated in 12-well plate on 18 mm glass coverslips (Karl Hecht, ref. 11817093) coated with 20 μg of human fibronectin (Merck, ref. FC010) in 1 mL of *regular culture medium* and allowed to attach for 48 hours. To eliminate estrogen activity, we prepared a culture medium aiming to minimize amounts of molecules with estrogen-like activity (i.e. phenol red and estrogens from the FBS). This medium, thereafter referred to as *estrogen-free medium*, is made of phenol red-free DMEM (Gibco, ref. 21063029) supplemented with 10% charcoal-stripped FBS (PanBiotech, ref. P30-2302). After the cells have attached to the coverslips in *regular culture medium*, cells were (step 1) washed twice in 1X PBS (Gibco, ref. 14190250) and once in phenol red-free DMEM; then (step 2) cultured in *estrogen-free medium* for two hours, with the medium replaced once during this period; then (step 3) placed in 1mL of *estrogen-free medium* containing 1 μM palbociclib (Sigma, ref. PZ0383) and cultured for 48 hrs. Finally (step 4) 1μL of DMSO containing 17-β-estradiol (E2, Sigma-Aldrich E2758) or not (vehicle control, labeled DMSO) was added 40 min before fixation. The final concentration of E2 in the medium was 100 nM.

Estrogen-deprived MCF7 cells are known to accumulate at the G1/S transition and to enter S-phase upon E2 induction^31,32^. To avoid confounding effect due to cell cycle progression in our study, we used palbociclib, a CDK4/6 inhibitor that blocks cells at the G1/S transition^33^, the efficacy of which we tested in our MCF7 cells by quantifying DNA synthesis by EdU incorporation^34^ over a period of 8 hrs, following 48 hrs in *estrogen-free medium* as in step 3 above but with or without 1 µM palbociclib and supplemented or not with 100 nM E2 (Figure S1C). This data confirms the effect of estrogen deprivation on cell cycle progression (marked increase of EdU-negative cells in estrogen deprived conditions) and the efficacy of palbociclib whether cells are estrogen deprived or not (0% of cells with EdU signal), consistent with previous reports^35^. Palbociclib was used as described above (steps 1–4) for all the data generated in this study, in which we observe induction of the studied estrogen-responsive genes (Figure 2C).

The experiments were performed on 5 different single-cell-derived populations of MCF7 cells and treated as replicates. To obtain the single-cell-derived populations, we separated individual cells from the population using flow cytometry and plated them individually in 96-well plates. Cells were then cultured in *regular culture medium* under standard conditions and upon reaching confluency replated to progressively larger culture dishes, until 15 cm plate. Cells were then collected and frozen at -80°C and transferred to liquid nitrogen for the long-term storage.

### RNA FISH probes

Custom RNA FISH probe sets of 20-nucleotide-long oligonucleotides were designed against *FOS, JDP2* and *BATF*, using the online Stellaris® Probe designer software (LGC, Biosearch Technologies) and ordered as DNA oligonucleotides conjugated respectively with CAL Fluor® Red 610, Quasar®670, and Quasar®570 fluorescent dyes, from Biosearch Technologies.

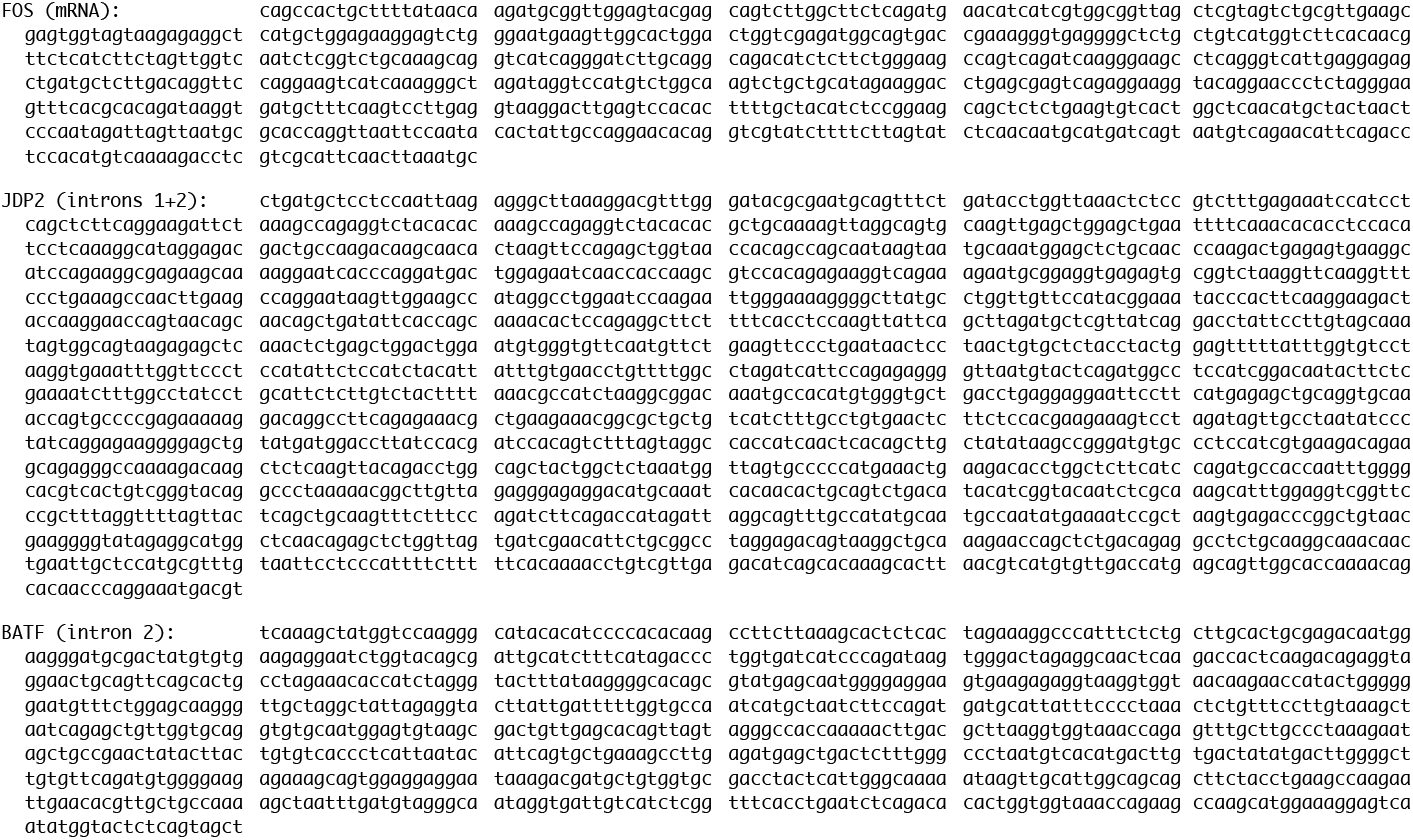

### Design and production of iFISH/Oligopaints library

A library of 12,005 oligonucleotides homogenously covering a 1.17 Mb genomic region encompassing our locus of interest (chr14:75,302,947-76,473,730, hg19 coordinates) was obtained from the ‘Full 40mers’ database of the iFISH platform^25^. To avoid the binding of DNA FISH probes to RNAs from our 3 genes of interest, all probes are (+)-strand sequences, hence targeting the (–)-strand, corresponding to the template (non-coding) strand of *FOS, JDP2* and *BATF*. The 40nt-long oligonucleotide sequences were extended according to Bintu *et al*.^36^ by adding (i) on either sides, 20nt-long sequences for secondary targeting (not used in this publication), then (ii) in 5’, a sequence (CATCAACGCCACGATCAGCT) to be targeted by a primer for the retro-transcription (RT) reaction, and finally (iii) on either sides, two sequences (CGGCTCGCAGCGTGTAAACG in 5’ and CGGTGCATTTGCGGGAAGAC in 3’) to be targeted by two PCR primers for amplification of the whole library. The resulting library of 140nt-long sequences was ordered from GenScript as a Classic Oligo Pool-12K oligonucleotide pool.

From the ordered oligonucleotide pool, DNA FISH probes were then produced as described in Beliveau *et al*.^37^ with small modifications in order to make the whole library of primary DNA FISH probes directly fluorescent. Briefly, the ordered oligonucleotide pool was first PCR amplified with Phusion Hot-start Master Mix (New England BioLabs), using forward primer CGGCTCGCAGCGTGTAAACG and reverse primer *TAATACGACTCACTATAGGG*GTCTTCCCGCAAATGCACCG, where the italicized part is a T7 promoter sequence. The PCR product was purified using DNA Clean & Concentrator-5 (DCC-25) kit (Zymo research). The resulting double-stranded DNA template was then *in vitro* transcribed using HiScribe T7 Quick High Yield RNA Synthesis Kit (New England BioLabs), producing a reverse-complement single-stranded RNA copy of the original library. This was then retro-transcribed with Maxima H Minus RT Transcriptase (ThermoFisher Scientific) using as RT primer a DNA oligonucleotide ATCGACCCGGCATCAACGCCACGATCAGCT ordered from IDT DNA coupled in 5’ to a 6-FAM (fluorescein) fluorophore. RNAs were finally degraded from the resulting RNA-DNA duplex using a NaOH-EDTA solution for 10’ at 95°C and the resulting fluorescently-labeled single-stranded DNA oligonucleotides were purified and concentrated using DNA Clean & Concentrator-25 (DCC-25) kit (Zymo research). All steps involving RNA were performed under RNase-free working conditions and with the addition of RNasin Plus RNase Inhibitor (Promega).

### Combined RNA-DNA FISH

The cells were crosslinked for 10 minutes at room temperature (RT) using Fixation Solution (4% paraformaldehyde (EMS, ref. 15710)/1X PBS (Gibco, ref. 14190250)) and subsequently washed twice for 10 min with 1X PBS. The cells were then permeabilized for 20 min at RT with Permeabilization Buffer (0.5% Triton X-100 (EUROMEDEX, ref. 2000-C)/1X PBS), washed three times for 1 min with 1X PBS and then treated with 0.1N HCl for 5 min. Next, the coverslips were incubated in the dark for 2 days in prehybridization buffer (2X SSC (EUROMEDEX, ref. EU0300-A)/50% formamide (Ambion, ref. AM9342)) at RT. After prehybridization, each coverslip was gently placed on microscope slides with a 30 μL drop of Primary Hybridization Buffer containing amplified primary iFISH/Oligopaints probes (2X SSC/50% formamide/10% dextran sulfate (Sigma, ref. D8906-50G)/0.2 μM of primary probes). The coverslips were then sealed with fixogum (MP Biomedicals, ref. 11430456) and left for 30 min in the dark at RT for the fixogum to solidify. Afterwards, the whole slides were placed on a heating block at 80°C for 3 minutes to denature DNA. Following denaturation, the slides were placed in a humid chamber and incubated overnight at 37°C in the dark. After hybridization of the primary probes, fixogum was gently removed and the coverslips were washed three times for 10 min with 2X SSCT (2xSSC/0.2% Tween) at 37°C, then twice for 7 min in 0.2X SSCT (0.2X SSC/0.2% Tween) at 65°C and finally briefly rinsed once with 4X SSCT (4X SSC/0.2% Tween) at RT and then twice with 2X SSC at RT. Subsequently, the coverslips were placed on microscope slides with a 25 μL drops of the Secondary Hybridization Buffer containing secondary smRNA FISH probes, as well as fluorescent readout probes aiming to visualize a small 20kb DNA region at the contact insulation site, although we did not use it in this publication (2X SSC/30% formamide/10% dextran sulfate/0.1 μM of each of the smRNA FISH probes/1 μM of secondary readout probes) in the same manner as for the primary hybridization. The coverslips were then sealed with fixogum, placed in a humid chamber and incubated overnight at 30°C in the dark. After secondary hybridization, the fixogum was gently removed and the coverslips were washed three times for 10 min in Wash Buffer (2X SSC/30%formamide) at 30°C, rinsed twice with 2xSSC at RT and then stained with 5 ng/mL DAPI in 2X SSC for 10 min at 30°C. The coverslips were then rinsed twice with 2X SSC, mounted on a microscope slide with Prolong Gold (Fisher Scientific, ref. P36934) and left overnight in the dark at RT to dry. After preparation, the slides were stored in the dark at -20°C.

### Microscopy and image quantifications

The FISH samples were imaged on an epifluorescence microscope made of a Nikon Ti-2 stand with a 60x 1.4NA CFI Plan Apochromat Lambda D oil immersion objective, a SpectraX (Lumencor) light source, a H117 XY stage (Prior), a Nano-ZL100 stage piezo (Mad City Labs), Semrock filters and dichroics, and an ORCA-Flash4.0 V3 sCMOS camera (Hamamatsu), and controlled by the MicroManager software^38^. Images were analyzed using custom-written analysis scripts (https://github.com/CoulonLab/FISHingRod). Nuclei were first segmented based on DAPI staining, using a threshold-based approach. Improperly segmented objects were excluded first based on their area, then through manual curation using only the DAPI channel. Fluorescent spots were detected in the DNA and RNA FISH channels. The 3D spot coordinates in each channel were corrected for chromatic shift using an affine transform (translation and rescaling) determined using images of broad-spectrum fluorescence beads (TetraSpeck, Invitrogen, ref. T14792) acquired on the same microscope.

Using, for the DNA FISH, a large library of 12,005 oligonucleotides that are directly fluorescent (no secondary oligonucleotide for labeling) permitted non-ambiguous identification of the alleles (fluorescence spot intensities of most alleles are well separated from background; Figure S1D). MCF7 cells having an instable karyotype, the copy number of alleles, even from a single-cell derived population, is variable (Figure S1E). We discarded nuclei with less than 2 alleles from our analysis, as well as nuclei with a high fluorescence background around RNA FISH spots. Plotting the 3D distances between the allele DNA loci and the detected RNA spots in each channel revealed a clear enrichment up to 650 nm (Figure S1F), suggesting that these correspond to nascent RNAs. Indeed, using this as a cutoff distance, RNA spots away from any allele loci showed a narrow distribution of fluorescence intensities, as expected for individual RNAs, while the fluorescence intensity distribution of RNA spots close to an allele locus showed a long tail towards larger values, as expected for genes with bursts of nascent RNAs (Figure S1G).

For each RNA channel, the single-RNA spot intensity was used to normalize fluorescence intensity values, yielding absolute numbers of RNAs. These normalization values showed little variation between experiments. When the spot intensity distribution lacked a clear single-RNA peak (e.g. due to high background), we used the value obtained in other experiments for the same channel. For each allele DNA locus, the amount of nascent RNAs was determined, for each channel, from the brightest RNA spot within 650 nm or assigned to 0 when no RNA spot was found within this distance (Figure 2C, top). Note that, as with most methods in the field, the resulting nascent RNA counts are underestimated, since RNAs become fully fluorescent only after the targeted region is fully transcribed and, for intronic probes, as long as it is unspliced. Furthermore, single nascent RNAs are indistinguishable from already-released single RNAs that happen to be within 650 nm of an allele DNA locus. Hence, we can only accurately determine the presence and number of nascent RNAs when the fluorescence signal exceeds a certain threshold (e.g. 2 single-RNA-equivalent fluorescence units for *FOS*; Figure S1G, dashed line). When binarizing our measurements to determine the fraction of active/bursting alleles (Figure 2C, bottom), we selected thresholds as low as possible while maintaining low false-positive calls: namely *c*_*FOS*_ = 2, *c*_*JDPS*_ = 2.5, and *c*_*BATF*_ = 2 in single-RNA-equivalent fluorescence units.

## Analysis of transcriptional correlations

### Population-level correlation patterns from public GRO-seq and Hi-C data

We used Global Run-On sequencing (GRO-seq) data set from Hah *et al*.22, mapping transcriptionally engaged RNA polymerases in MCF7 cells upon induction with 100 nM estradiol (E2). From the provided table (Gene Expression Omnibus GSE27463) quantifying GRO-seq reads in all RefSeq-annotated genes, we note *𝒳*_*k*_ and 𝒴_*k*_ the measurements obtained on two genes X and Y in experimental condition *k* (i.e. at 0’, 10’, 40’ or 160’ after induction with E2) and *𝒳* = ∑_*k*_ *𝒳*_*k*_⁄𝒩 and 𝒴 = ∑_*k*_ 𝒴_*k*_⁄𝒩 their means, where 𝒩 = 4 is the number of experimental conditions.

The covariance and Pearson correlation, across experimental conditions, between the population-level measurement of the transcriptional activities of genes X and Y are

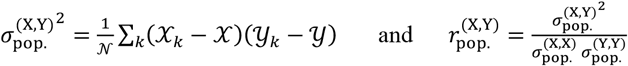

This correlation measures how similar the *shapes* of the time profiles of the transcriptional response of two genes are, up to *arbitrary offset* and *rescaling* of either time profile —that is, independently of basal transcription levels and of the magnitude of the E2-induced response. In other words, the correlation between (𝒳_0′_, 𝒳_10′_, 𝒳_40′_, 𝒳_160′_) and (𝒴_0′_, 𝒴_10′_, 𝒴_40′_, 𝒴_160′_) is the same as between (*a*𝒳_0′_ + *b,a*𝒳_10′_ + *b,a*𝒳_40′_ + *b,a*𝒳_160′_ + *b*) and (𝒴_0′_, 𝒴_10′_, 𝒴_40′_, 𝒴_160′_)for any value of *a* > 0 and *b*.

Pearson correlation values close to 1 indicate similar transcriptional time profiles (e.g. the activity of two putative genes A and B both increasing after 10’, plateauing until 40’, then reverting at 160’, even if A starts with a high basal level and shows small E2-induced variations while B does not). Values close to 0 indicate no similarity (e.g. gene A *vs* a gene C which activity keeps increasing). Values close to -1 indicate opposite transcriptional time profiles (e.g. gene A *vs* a gene D which would be downregulated after 10’ and until 40’, then reverting at 160’).

Figure 1A presents the 2D histogram, for all gene pairs genome-wide within 1 Mb of each other, of the Pearson correlation between their transcriptional response to E2 and the genomic distance between their promoters. Figure 1B presents the same analysis but only for the top 20% most insulated gene pairs, as calculated from the Hi-C data of Rodriguez *et al*.23 obtained in MCF7 cells (Gene Expression Omnibus GSE121443). For this, we used the cooler package^39^ to generate 1 kb- and 10 kb-resolution matrices for each experiment, pooled them together, and calculated an insulation score at each genomic position by summing the contact frequency values observed between two 140kb-regions upstream and downstream of the position of interest, leaving a 10kb-gap between them. A *low value* of the insulation score indicates a *strong contact insulation*.

### Approach 1 – Separating *cis* (allele-intrinsic) and *trans* (allele-extrinsic) components in transcriptional correlations from RNA FISH data

#### – Principle –

Consider we have nascent RNA measurements at a total of *n* alleles from a given experiment. Let (*x*_*i*_,*y*_*i*_) be the *n* measurements from genes X and Y at each individual allele *i*, and their means be *x* = ∑_*i*_ *x*_*i*_⁄*n* and *y* = ∑_*i*_ *y*_*i*_⁄*n* (unless otherwise noted, sums on *i* are taken over [1,*n*]).

The covariance across alleles of the allele-level measurements is

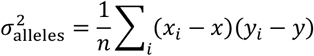

Let us consider the *n* alleles to be partitioned among *N* cells. Let *n*_*j*_ denote the number of alleles in cell *j*, so that *n* = ∑_*j*_ *n*_*j*_ (sums on *j* are taken over [1,*N*]). Let us note the cell-wise summed measurements as 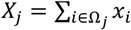 and 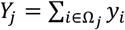 where *i* ∈ Ω_*j*_ denotes the alleles *i* in cell *j*. Using the same mean subtraction as above, such that 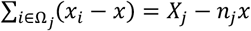 and 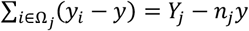, the covariance of the cell-wise summed measurements is

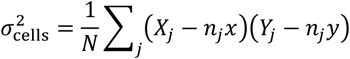

We note that 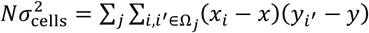 enumerates all the possible pairs within cells, including both *cis* and *trans* pairs. Hence, the covariance over all possible *trans* pairs, i.e. all the possible intra-cellular permutations (explicitly 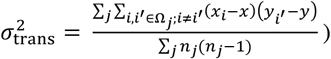) can be simply calculated as

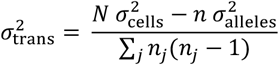

as illustrated in Figure S2A.

The part of 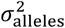 explained by cell-wise (allele-extrinsic) effects corresponds to 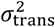 and their difference, which we note 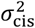, isolates the part of the covariance originating from local (allele-intrinsic) effects *in cis*:

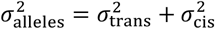

If the covariance across alleles were fully explained by *trans* effects, then 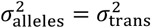 and 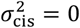 (Figure S2B, left). On the opposite, if alleles behaved independently and there was no cell-to-cell heterogeneity, then 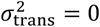 and the measured covariance would fully come from *cis* effects 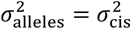 (Figure S2B, right). In the general case, 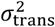 quantifies cell-to-cell heterogeneities or other sources of inter-allelic coupling and 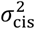 is what would be observed if there were no such inter-allelic effects.

#### – Remarks –

1. In the case of two identical copies of a gene in each cell, our approach simplifies into the classical *extrinsic* and *intrinsic noise* definition^26^, which decomposes the variance as 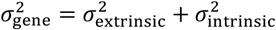. Denoting 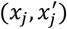 the two measurements in each cell *j*, we have 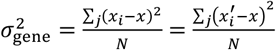 and 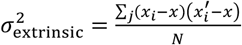 with 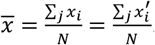
2. We chose to use cell-wise summed measurements (*X*_*j*_,*Y*_*j*_), but this approach can be adapted to use cell-wise averages 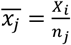 and 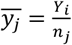. Namely, we write

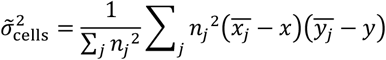

(notice the unusual weighting) which only differs from 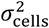 by a scaling factor corresponding to the average number of allele pairs per cell (i.e.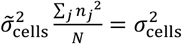). Then, we get 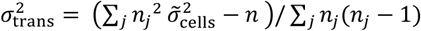 and the rest is identical.
3. Although related, this approach differs from the classical way to separate within-group and between-group effects using the *law of total covariance*, which reads:

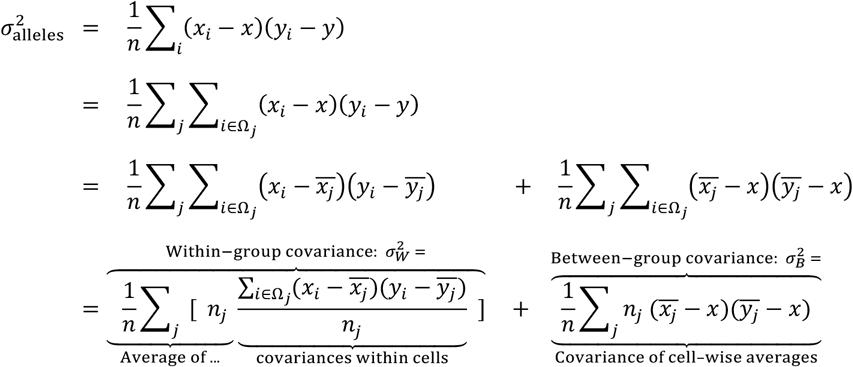

Notice how the weighting of the terms in the sum differs between 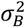 and 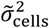 above. Substituting 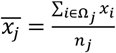 and 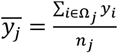 leads to 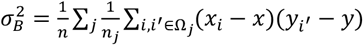 and 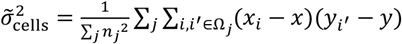. Both expressions enumerate all the possible intra-cellular pairs, but the former gives them unequal weighting between cells, making it impossible to subtract the *cis* pairs using 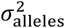.

Two useful limit cases to consider are:

- If all cells have the same number of alleles, then 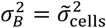. In this case, we get

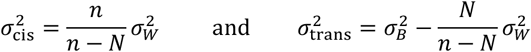
- In addition, for large groups (*n*/*N* ≫ 1, which does not hold here for alleles in cells, but may be valid in other situations; see *Generalization* below), we get

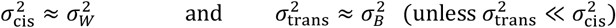

#### – Application –

Applying this approach to different gene pairs, e.g. X, Y, Z, yields covariance matrices:

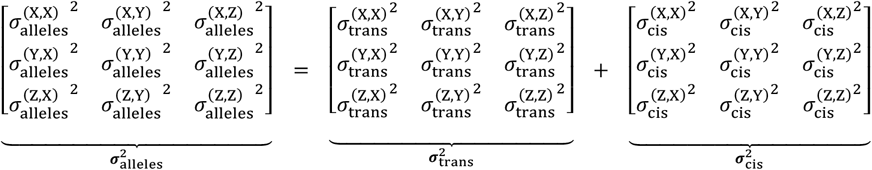

Figure S3 shows the values of covariance matrices 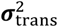 and 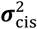 built from the nascent RNA FISH data on *FOS, JDP2* and *BATF*, in both uninduced (DMSO) and induced (E2) experimental conditions. For each gene and each condition, the *cis*-to-*trans* ratio is displayed (e.g. in E2 condition, 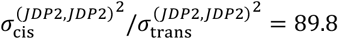), as well as the percentage of its *cis* variance that is explained by its covariance with either of the other two genes (e.g. in E2 condition, 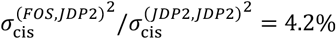 and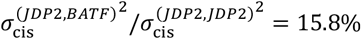).

The Pearson correlation across alleles, between any gene pair X and Y, can be decomposed as

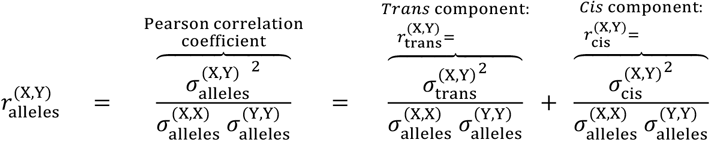

which is what Figure 3A represents.

Note that the denominators are the same so that 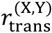 and 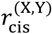 represent two additive components of 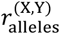. Hence, 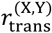 and 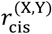 are <u>not</u> the Pearson correlation coefficients of the *cis* and *trans* components (i.e. calculated from the covariance matrices 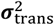and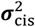). However, since most of the variability occurs *in cis* in our case (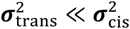, Figure S3B), the *cis* and *total* covariances are very similar 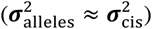, so that what is shown on Figure 3A (i.e. the *cis* component of the Pearson correlation, 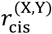 above) is very close to the Pearson correlation of the *cis* component of the variability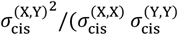).

#### Generalization

Our approach can be applied recursively to experimental data where sample elements are organized hierarchically. In such cases, the approach is applied iteratively, starting from the lowest level, and each iteration uses the aggregated data from the previous iteration as its new elementary data.

As illustrated in Figure S2C, after applying our approach to nascent RNA counts from individual alleles (level 0) and their cell-wise aggregated values (level 1; i.e. summed or averaged within each cell), if the data came for instance from a spatial transcriptomics experiment on a tissue section with various cell types and tissue areas, a second iteration could be applied to the cell-level measurements (level 1) and their aggregated values over cells of the same cell type (level 2). A third iteration could then be applied to cell-type-level measurements (level 2) and their aggregated values over various tissue areas (level 3), and so on.

Beyond this example, our approach is not limited to FISH-based transcriptomics data. It may also be well-suited to analyze data from single-cell and spatial multi-omics technologies^40^, including both microscopy- and sequencing-based methods, since the resulting data is often intrinsically multi-level and hierarchical. (Note that allele-level measurements are not always possible with these approaches, in which case the lowest level in the hierarchy can be directly the single-cell level). In this context (Figure S2D), our approach can be readily extended by incorporating:

i. at the lower level in the data hierarchy: alongside or instead of gene transcription, other single-cell ‘omics’ measurements, such as DNA accessibility by single-cell ATAC-seq, or protein occupancy or histone marks by single-cell CUT&Tag;
ii. as extra higher levels in the hierarchy: beyond spatial organization (groups and subgroups of cells, as in Figure S2C), temporal aspects (e.g. developmental time in an organism, response to treatment or perturbation) and experimental conditions (e.g. drug treatments).

### Approach 2 – Separating the contributions of burst co-occurrence and burst size correlations

#### – Principle –

Consider the contingency table obtained by classifying genes X and Y at each allele as undergoing or not a transcriptional burst. This equates to binarizing the data into 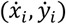 as

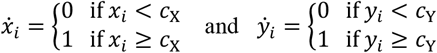

where (*c*_X_,*c*_Y_) are the thresholds to consider the genes as undergoing a transcriptional burst. The associated contingency table is

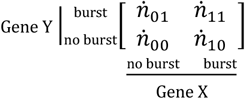

where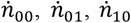, and 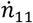 sdenote the number of alleles where 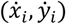 are respectively (0,0), (0,1), (1,0), and (1,1) (with 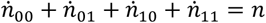), corresponding to the partition 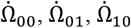, and 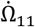 of the ensemble of alleles Ω.

This contingency table describes the correlations of the burst occurrences between the two genes, irrespective of burst sizes. The associated covariance is

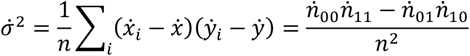

with 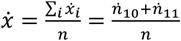 and 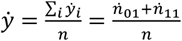 the fraction of bursting alleles for either gene.

What follows shows how the covariance 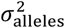 of the nascent RNA counts (*x*_*i*_,*y*_*i*_) can be decomposed as a weighted sum of a contribution from the covariance of the binarized data 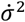, **describing correlations of burst occurrences** independently of their sizes, and other components, altogether noted 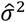, **describing correlations involving burst sizes**.

We first describe this approach independently of the previous one to separate *cis* and *trans* effect and address in the subsequent subsection how both approaches can be combined. For consistency, we first note here 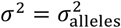 the covariance of the non-binarized data (*x*_*i*_,*y*_*i*_).

#### – Definitions and notation –

Let us introduce the following terminology and notation:

- We call *quadrants* the four regions of the (*x*_*i*_,*y*_*i*_) scatter plot, defined by the thresholds *c*_0_ and *c*_2_, corresponding to the partition of alleles into 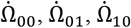, and 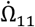.
- *Quadrant-wise averages*: We note 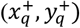 the averages of (*x*_*i*_,*y*_*i*_) within each quadrant

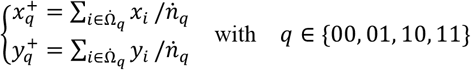

and use the shorthand 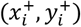 to represent, for each allele *i*, the values of 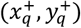 in its quadrant (i.e. 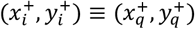 with *q* such that 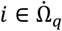). The notation ‘+’ symbolizes the (*x*_*i*_,*y*_*i*_) scatter plot being divided into quadrants.
- *Half-plane marginal averages* are defined as follows:
  ∘ 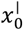 and 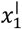 denote the averages of the *x*_*i*_ values that are respectively lower or greater than the threshold *c*_X_

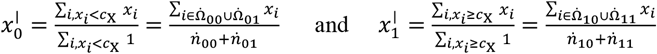
  ∘ and 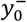 and 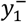 are the averages of *y*_*i*_ that are respectively lower or greater than *c*_Y_

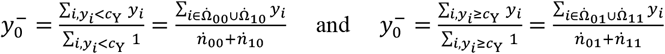

These are *marginal* quantities since 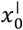 and 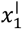 are defined independently of *y*_*i*_, and 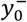 and 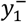 independently of *x*_*i*_. On the (*x*_*i*_,*y*_*i*_) scatter plot, 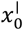 and 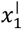 (resp. 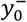 and 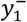) are the averages within the two *half-planes* defined the vertical (resp. horizontal) threshold *c*_0_ (resp. *c*_2_). As a notation shorthand, the value 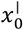 or 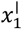 (resp. 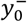 or 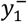) corresponding to each allele *i* and each quadrant *q* are noted 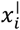 (resp.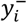) and 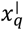 (resp.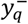).
- *The binarized data* (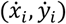), as defined earlier, are 0 and 1 values representing whether *x*_*i*_ (resp. *y*_*i*_) is below (0) or above (1) the threshold *c*_X_ (resp. *c*_Y_).

To give a few concrete examples: 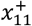 is the average number of nascent RNAs on gene X when both genes are bursting simultaneously. 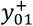 is the average number of nascent RNAs on gene Y when gene Y is bursting and gene X is not. 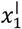 is average number of nascent RNAs on gene X when bursting. 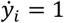 indicates that gene Y is bursting on allele *i*, in which case 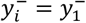.

Note that the raw nascent RNA counts (*x*_*i*_,*y*_*i*_), their quadrant-wise averages 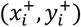, and their half-plane marginal averages 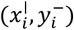 all have the same average (*x,y*) 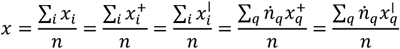 and 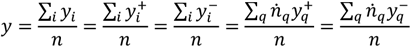 and that 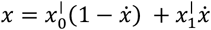 and 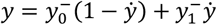.

#### –Derivations –

Our approach to connect the statistics of the RNA counts and the binarized data consists in expressing the former from the latter, using the three following steps:

i. deviation from quadrant-wise averages
ii. deviation from half-plane marginal averages
iii. affine transform of binarized data (same as above)

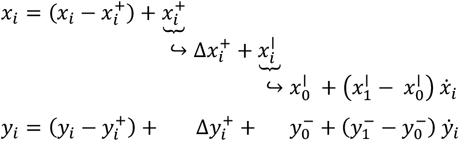

and examining how this breaks down the covariance into meaningful additive components. Step (i) consists in applying the *law of total covariance* to the four quadrants:

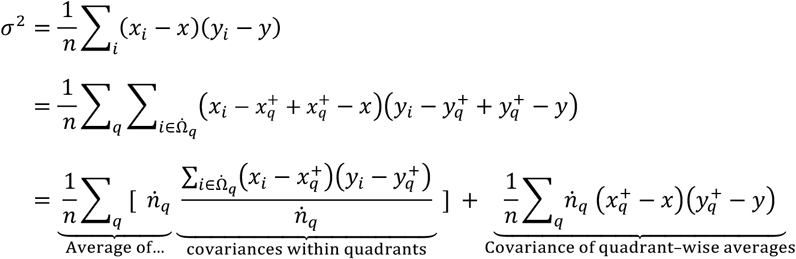

The first term quantifies the covariance within each individual quadrant, among which quadrant 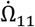 describes how burst sizes are covary when both genes are bursting simultaneously, while the other quadrants should have a small contribution since one or both genes have close to zero nascent RNA count.

The second term quantifies how the average nascent RNA counts within quadrants covary across the four quadrants. It differs from the covariance 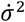 of the contingency table because, although the number of elements in each quadrant are the same as in the contingency table 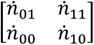, the associated values are not the *unit square* 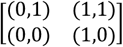, but 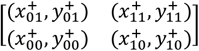,which in the general case is a non-rectangle quadrilateral. If it were a rectangle, then, as we show below, this second term in the equation above would simply be 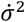 multiplied by a prefactor. Hence, the rational for steps (ii) and (iii) is to map this quadrilateral first onto a rectangle, then onto the unit square.

Step (ii) consists in re-expressing the quadrant-wise averages 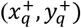 in terms of deviations 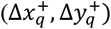 from the half-plane marginal averages, noted 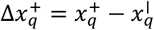 and 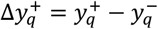.

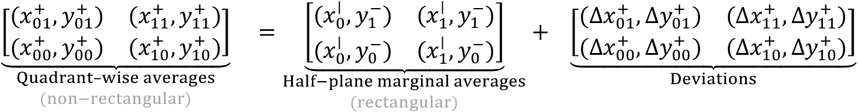

The deviations terms 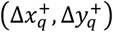 capture how the average burst size of either gene varies when conditioned on the fact that the other gene is bursting or not. For instance, 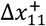 and 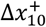 represent by how much the average nascent RNA counts on gene X during a burst (i.e. 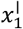) changes knowing whether gene Y is bursting or not (i.e. difference with 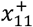 or 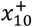).

The covariance of the quadrant-wise averages (second term in the previous equation) rewrites

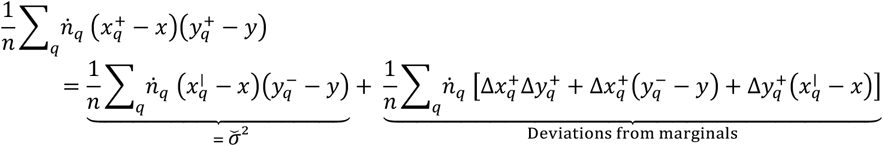

where we factored out into the second term all the contributions of the deviation 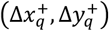. This second term hence represents how the size of bursts on either gene depends on the presence of a burst on the other gene. Combined with the average covariance within quadrants (first term of the previous equation), this leads to a term we call 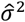, capturing altogether how the size of bursts on either gene depends on the activity of the other gene:

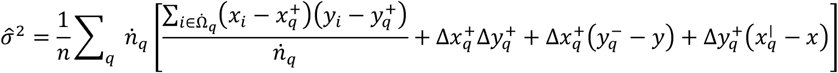

Finally, the remaining term, labeled 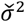 above, is a simple rescaling of 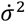, the covariance of the contingency table. Indeed, applying step (iii) to express by rewriting and substituting the means (*x,y*) by their expression in terms of 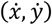, we obtain

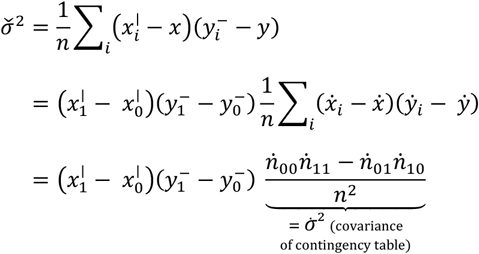

Therefore, we obtain

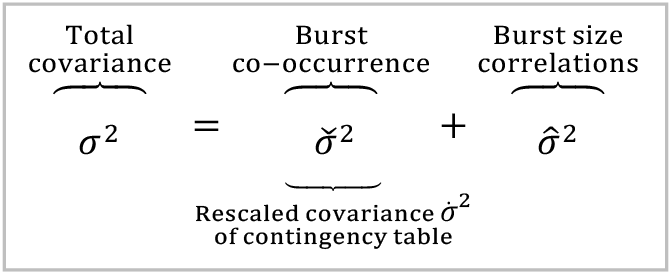

The first term on the right-hand side stems from the contingency table and exclusively **quantifies how the occurrence of bursts correlates between genes**, irrespective of their sizes. The second term captures effects related to burst sizes, **quantifying how the size of bursts on either gene correlates with the activity of the other**.

#### –Application–

Practically,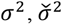, and 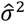 can be easily calculated as follows:

a. Calculate the covariance *σ*^2^ from the nascent RNA counts (*x*_*i*_,*y*_*i*_).
b. Calculate the covariance 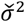 from the 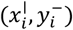 values (simply obtained by replacing all the *x*_*i*_ values below *c*_X_ by their average, all the *x*_*i*_ values above *c*_X_ by their average, all the *y*_*i*_ values below *c*_Y_ by their average, all the *y*_*i*_ values above *c*_Y_ by their average).
c. Calculate the third covariance as the difference between the first two: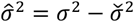.

Correlation matrices are built by considering different gene pairs for X, Y, … etc.

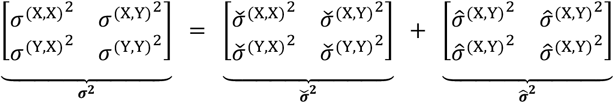

Using this decomposition of the covariances, the Pearson correlation can be separated into:

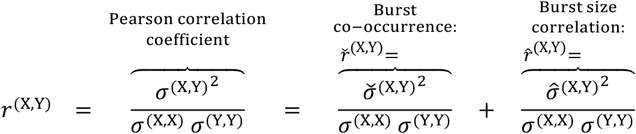

Note that the denominators are the same, so that *ř*^(X,Y)^ and 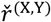 represent two additive components of *r*^(X,Y)^. Hence, 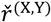 and *ř*^(X,Y)^ are <u>not</u> the Pearson correlation coefficients calculated from the covariance matrices 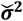 and 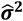.

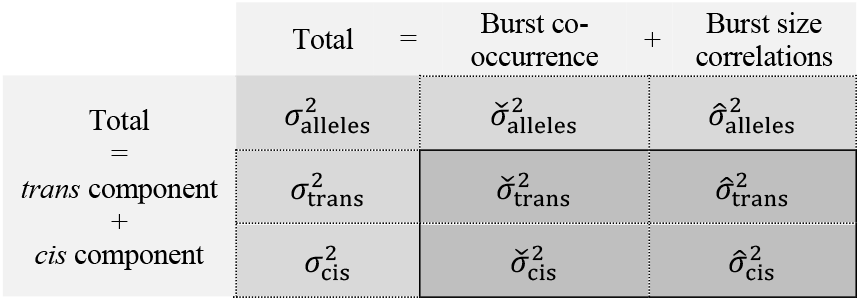

#### Combining both approaches

Our two approaches to separate (i) *cis* and *trans* contributions and (ii) burst co-occurrence and burst size correlation effects are orthogonal and can be combined. Indeed, the former holds for any allele-level measurements grouped by cells. Hence, we use the exact same equations (page 16) and same categorization of the *n* alleles into *N* cells (i.e. ensembles Ω_*j*_) to

a. calculate 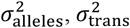, and 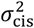 from the (*x*_*i*_, *y*_*i*_) values and their cell-wise sums,
b. calculate 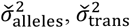, and 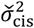 from the 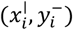 values and their cell-wise sums.

Then, the two approaches are orthogonal in the sense that it is equivalent to:

(c.) apply the equations on page 16 to obtain 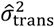, and 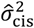 from the allele-level and cell-level covariances calculated as 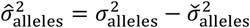
c. or, more simply, deduce directly all three covariances 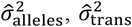, and 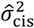 by subtracting 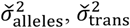, and 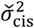 from 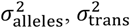 and 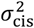.

Figure 3B represents the separation of 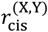 into the two additive sub-components 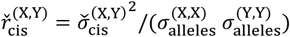 and 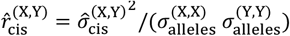. Denominators are allele-level standard deviations, since 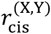 is itself already a component of 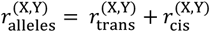.

#### Sample sizes, replicates, and statistical analysis

Both DMSO-control and E2-induced experiments were performed in 5 replicates, using 5 different single-cell-derived MCF7 cell populations (see *Cell lines and cell culture*). The number of analyzed alleles (*n*) and nuclei (*N*) for all 5 replicates were *n* = 6093, 17066, 12637, 13255, 11148 and *N* = 1160, 2890, 2187, 1835, 1869 in DMSO condition and *n* = 7005, 7157, 16476, 21658, 8647 and *N* = 1274, 1523, 2649, 3159, 1439 in E2 condition. Pearson correlation coefficients and their sub-components were calculated independently for each replicate and all statistical tests were then performed across the 5 replicates using either a 2-sample two-sided t-test with Welch correction (to account for unequal variances) or a 1-sample two-sided t-test, as reported in the following table.

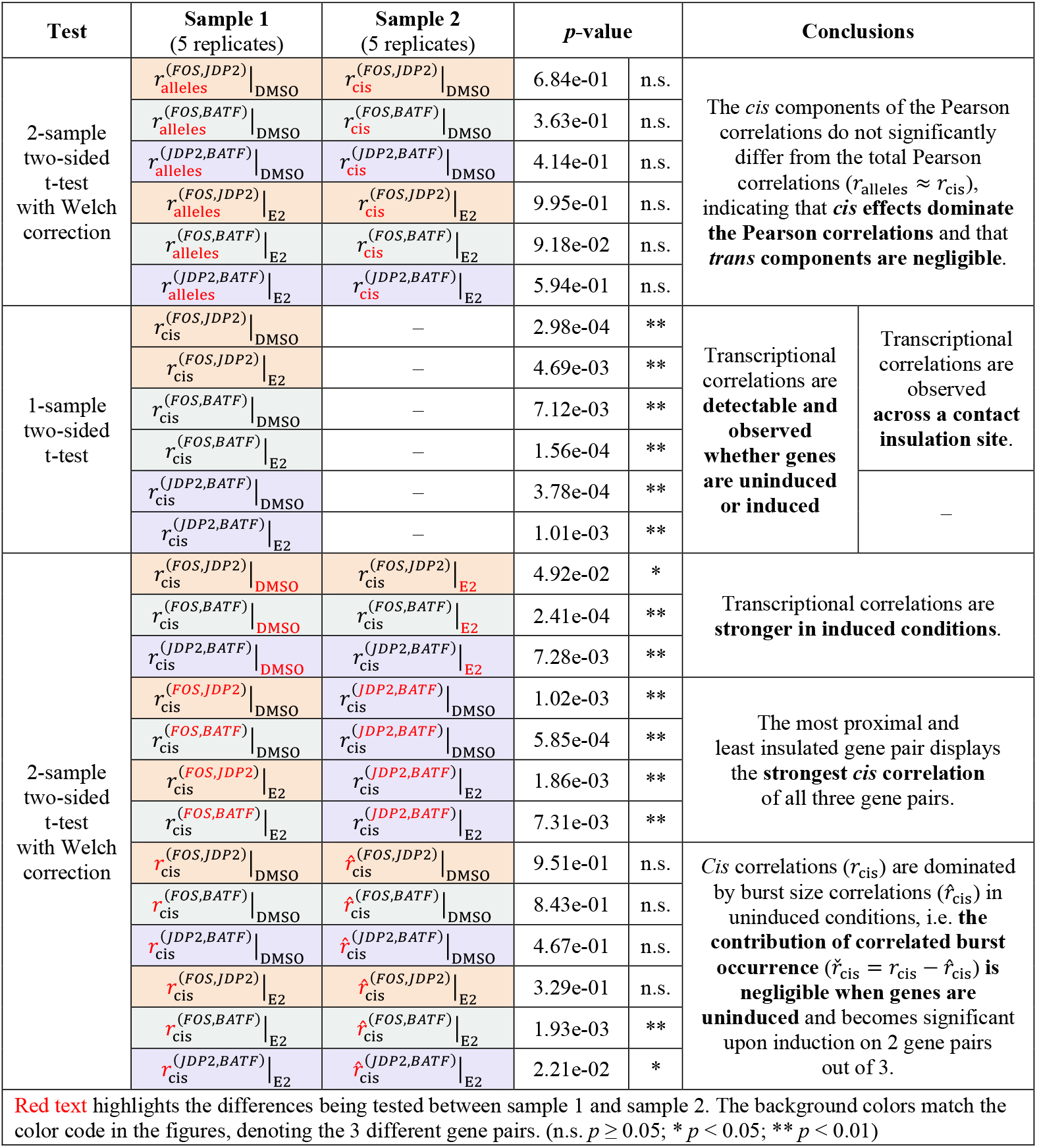

#### Resource availability

Data and code are available on a Zenodo repository at http://doi.org/10.5281/zenodo.14865160. This includes all the processed data and analysis scripts, and a subset of raw image files (i.e. first images for each replicate) to match the size limitations of Zenodo repositories (50 GB). The whole dataset (1.6 TB) is available upon request. Image analysis scripts for spot detection and nuclear segmentation are available at https://github.com/CoulonLab/FISHingRod.

## Acknowledgments

The authors wish to thank Magda Bienko for advice on FISH approaches, Vittore F. Scolari for help with Hi-C data processing, and all the members of our research group for discussions and comments. We also thank the Flow Cytometry Core Facility of Institut Curie (CYTPIC), the BMBC platform of UMR168, and the Cell and Tissue Imaging Facility (PICT-IBiSA) of Institut Curie, member of the national infrastructure France-BioImaging (https://ror.org/01y7vt929) supported by the French National Research Agency (ANR-24-INBS-0005 FBI BIOGEN). This work received funding from the Centre National de la Recherche Scientifique (CNRS); the Institut Curie; the ATIP-Avenir program of CNRS and INSERM, the Plan Cancer of the French ministry for research and health; the European Union’s Horizon 2020 research and innovation program, through the Marie Sklodowska-Curie grant agreement No 666003 (H2020-MSCA-COFUND-2014 IC-3i PhD program of Institut Curie) and the European Research Council (ERC) (grant agreement 757956, project “4D-GenEx”); the LabEx DEEP (ANR-11-LABX-0044 and ANR-10-IDEX-0001-02); and the program Fondation ARC (grant agreement PJA 20161204869).

## Author contributions

MAK, KJEB, and AC designed the study. AC obtained funding. MAK and KJEB performed experiments. IAL, JR, CJ, and AB produced essential reagents. MAK, KJEB, and AC developed analysis methods and performed data analysis. MAK, KJEB, and AC wrote the manuscript with inputs from all the co-authors.

## Declaration of generative AI and AI-assisted technologies

Generative AI was used occasionally during the preparation of this publication for enhancing clarity. After using this tool, the authors reviewed and edited the resulting text as needed and take full responsibility for the content of the publication.

